# Endogenous short enhancer sequences increase expression of soybean and cowpea RUBP regeneration genes

**DOI:** 10.64898/2026.04.29.721404

**Authors:** Micha Wijesingha Ahchige, Virginie Mengin, Christine A. Raines

## Abstract

Improving regeneration of ribulose-1,5-bisphosphate (RUBP) is a promising approach to improve photosynthesis and plant growth. In addition to transgenic overexpression of target genes, it could be possible to directly overexpress endogenous target genes, through transcriptional enhancements. As shown by the recent discovery of a short sequence motif, that resembles the known octopine synthase (ocs) enhancer, transcriptional enhancement is achievable by relatively short endogenous sequences. In this study, we query the genome of several model and crop plant genomes for the presence of short enhancer motifs. We find hits across all genomes including some in promoter regions of genes. By using derivatives of these motifs in a transient fluorescence assay, we show that several of these are capable of inducing target gene expression in different promoter contexts. A motif scan of the created constructs, for the presence of known transcription factor binding sites, shows that the insertion of these motifs has created binding sites for different TGA-, NAC- and bZIP-transcription factors. Taken together our study shows the feasibility of finding enhancer sequences in the genomes of different plants. With advancement in gene-editing technologies, like prime editing, using such endogenous enhancer sequences, could allow for precise cisgenic promoter engineering of target genes.

## Introduction

It is currently expected, that the rising demand for crop production by 2050, will not be met by present trends in yield increase of key global crops [1]. As such it will be necessary to further improve the yield increases through different strategies. Different approaches to improve photosynthetic efficiency have been suggested, that could collectively be able to double the yield potential of major crops [2]. As one such strategy, optimizing the regeneration of ribulose-1,5-bisphosphate (RUBP) has already been shown to be able to increase assimilation and growth of tobacco plants in the greenhouse as well as in field experiments [3, 4]. RUBP acts as the acceptor molecule for CO_2_ and it takes 8 enzymes to regenerate it [5]. One of the enzymes of RUBP regeneration is Sedoheptulose-1,7-bisphosphatase (SBPase) and a modelling approach could show that allocating more resources to this enzyme could improve overall assimilation rate [6]. By means of transgenic overexpression of this enzyme, it could be shown that this can indeed improve overall assimilation, as well as plant growth and biomass [3, 4]. Building on these conventional transgenic approaches we were also interested to develop advanced gene-editing approaches to be able to directly overexpress the endogenous enzymes in target crops. A recent study has found a 12 bp palindromic sequence, called plant enhancer (PE), in the maize genome and the insertion of this sequence as a triplicate (3xPE) into the promoter of target genes can significantly increase their gene expression [7]. The discovered element is identical to the core of a 16 bp palindrome from the promoter of the octopine synthase (ocs) gene of *Agrobacterium tumefaciens*, that has been shown to function as an enhancer and was named OCS element [8]. A saturation mutagenesis of this element suggested that some enhancing capability can be retained even with most single and double base substitutions but strongly declines with more base changes and single deletions [9]. Similar elements have been found in the promoter regions of other T-DNA genes and plant viral promoters and a 20 bp consensus sequence has been built between them [10]. Gel retardation assays in these studies showed two retarded bands, which suggested the binding of a protein factor, named OCSTF, as a monomer or dimer to the sequence [9, 10]. It could further be shown that the presence of the upper band, i.e. the dimer, is linked to transcriptional activation [9]. Following these early studies, several other OCS binding factors (OBF) have been found, which are now mostly referred to as TGA transcription factors [11–14]. The recent discovery of the plant enhancer has demonstrated great potential of OCS-derived motifs for the overexpression of target genes [7], which has prompted us to explore this approach to find additional motifs and investigate the potentially binding factors.

In this study we query the genomes of different model and crop plants for the presence of OCS-like motifs. We then insert the putative enhancers into promoters of our target genes to drive the expression of YFP and measure fluorescence intensity in a transient *Agrobacterium*-tobacco leaf infiltration assay. Finally, we scan the created promoter for potential binding sites of known transcription factors. Our work shows several motifs that can induce target gene expression and thus highlight an approach to find endogenous enhancer motifs that could be used for cisgenic-gene editing of target crops.

## Material and Methods

### Constructs and cloning

Genomic DNA was isolated from soybean (*Glycine max* cv. LD11) and cowpea (*Vigna unguiculata* IT86D-1010) with the DNeasy® Plant Pro Kit (Qiagen) according to manufacturer’s instructions. Primers were designed to amplify the 1000 bp upstream of the first codon, which includes the 5’UTR and the promoter. Promoter constructs were generated by first amplifying genomic DNA via regular PCR and then introducing different sequences via overhang PCR using VeriFi® Hot Start Mix (PCR biosystems) or Q5® High-Fidelity DNA Polymerase (NEB)(see Supporting Data S 1 for a list of all used primers). The custom level 2 double fluorophore vector was generated by assembling a lacZ fragment, a fragment containing the EYFP coding sequence and an HSPt-terminator together with a fragment containing a p35S-promoter, the coding sequence of mRFP1 and a NOSt-terminator, together with a linker into pAGM4723 via Golden Gate cloning using BbsI-HF® (NEB) and T4 DNA ligase (NEB), according to previously published protocols [15, 16]. The fragments originally derived from pICH47732, pICSL80014, EC15320 and pICH41766 and the mRFP1 expression cassette was ordered from Twist Biosciences (Twist Biosciences). The plasmids pICH47732, pICSL80014, pICH41766 and pAGM4723 are derived of the MoClo toolkits[17]. Promoter fragments were then introduced upstream of EYFP into the custom vector via the same Golden Gate protocols described above but using BsaI-HF®v2 (NEB) as the restriction enzyme. From the Golden Gate reaction 5 µL were used for transformation of 25 µL NEB® 10-beta Competent E. Coli cells (NEB). Individual colonies were picked and tested via colony PCR using Thermo Scientific™ DreamTaq™ Green DNA Polymerase (Thermo Scientific). Plasmid DNA was isolated from putatively positive clones using NucleoSpin® Plasmid kit (Macherey-Nagel) and sent for Sanger sequencing (Eurofins). Sequencing files were aligned to genome files of the target crop cultivars *Glycine max* LD11 (PRJNA1137955) and *Vigna unguiculata* IT86D-1010 [18]. Correctly assembled plasmids were transformed via electroporation into electrocompetent *A. tumefaciens* GV3101 by adding 500 ng vector DNA to 40 µL cells in electroporation cuvettes (Molecular BioProducts) with a 2 mm gap and using an eporator (Eppendorf) at 2500 V. Individual cells were picked and presence of the plasmid was again confirmed with another round of colony PCR as described above.

### Plant cultivation

Tobacco plants were cultivated in a phytotron (fitotron, Weisstechnik,) by sowing seeds of *Nicotiana tabacum* cv. Samsun directly onto moist soil (Levington® Advance Seed and Modular + Sand F2 + S, ICL). After 6-8 days individual seedlings were transplanted into 3-inch pots with moist soil (Levington® Advance Seed and Modular + Sand F2 + S, ICL) and kept in trays. Plants were watered 3 times per week. Throughout the entire growth period, plants were kept in a (14h/10h) light-dark cycle, with temperatures of (26°C/26°C), humidity of (65%/65%) and light intensity of (∼200 µmol*m^-2^*s^-1^/0 µmol*m^-2^*s^-1^). Plants were ready to be used for infiltration assays, 5 weeks after sowing seeds.

### Agro infiltration

From each 5-week old plant two mature leaves were used for infiltration with transformed *A. tumefaciens* according to an established infiltration protocol [19] with some minor modifications. The modifications comprised using Na_2_HPO_4_ instead of Na_3_PO_4_ and using 100 µL 50mM acetosyringone instead of 5 µL 1M acetosyringone for the infiltration medium. Plants were thoroughly watered the day before the assay to ensure widely opened stomata. Per assay, 8 plants were used and constructs were infiltrated onto 4-8 infiltration spots per leaf. Per leaf one spot was infiltrated with wild type *A. tumefaciens* without any vector as a blank to account for background fluorescence noise.

### Fluorescence measurement

After infiltration, plants were kept for 3 more days in the phytotron. From each infiltration spot, 3 leaf discs with 6 mm diameter were punched out using a Unicore Punch (Qiagen) and transferred to Thermo Scientific™ Black 96-Well Immuno Plates (Thermo Scientific), containing 400 µL milliQ water with the abaxial side facing up, similar to an established protocol [20]. YFP and RFP fluorescence was measured with a CLARIOstar Plus plate reader (BMG Labtech). YFP was excited at 497 nm with a bandwidth of 15 nm and measured at 540 nm with a bandwidth of 20 nm and RFP was excited at 584 nm with a bandwidth of 15 nm and measured at 626.5 nm with a bandwidth of 20 nm. Leaf discs of the same construct across the two leaves of the same plant were considered technical replicates and plants were considered biological replicates. Per plate the median fluorescence value of leaf discs from blank infiltrated regions was calculated. Individual leaf discs from spots infiltrated with the blank that deviated more than 0.5x from this value were excluded from further normalization. Per leaf the average blank fluorescence intensity value was subtracted from the fluorescence intensity of all leaf discs. For each leaf disc a YFP/RFP ratio was calculated and then the average was calculated for leaf discs from two leaves from the same plant. The resulting values were treated as biological replicates and used for downstream statistical analysis.

### Computational analysis and software

Phytozome [21] and Solgenomics [22] were used to obtain whole genome data of *Glycine max* Wm82.a4.v1 [23], *Vigna unguiculata* Vunguiculata v1.2 [24], *Manihot esculenta* Mesculenta v8.1 [25], *Zea mays* Zmays RefGen V4 [26], *Oryza sativa* Osativa v7.0 [27], *Nicotiana tabacum* Nitab v4.5 [28], *Arabidopsis thaliana* Athaliana Araport11 [29], *Solanum lycopersicum* Slycopersicum ITAG 3.2 [30], *Hordeum vulgare* Hvulgare Morex V3 [31] and *Triticum aestivum* Taestivum cv Chinese Spring v2.1 [32]. Based on these genomes an artificial genome was simulated. This was achieved by taking the median number of chromosomes and distributing the median genome size equally across the chromosomes. The artificial genome was then created by randomly sampling the nucleotides “A”, “C”, “G” and “T” and concatenating them to form artificial chromosomes. Manual sequence manipulation and in silico cloning was performed with Benchling [33]. Primers were designed with Primer-BLAST and Benchling [33, 34].

The majority of computational analysis and visualization was performed using R statistical software [35] version 4.4.0 in the RStudio IDE [36] version 2023.12.1.402. Genomes were queried for the presence of small enhancer sequences using NCBI BLAST [37] on a high performance computing cluster and using the R package Biostrings [38]. For the NCBI BLAST search, the query sequence was the 16 bp OCS sequence described before [8] and the short sequences parameter was used. For the Biostrings search the query sequence was the 20 bp OCS consensus sequence described before [10] and up to two mismatches were allowed, while the subject sequence was fixed. Hits were considered to occur in the promoter of genes if they fell at least partially, into the sequence 1 kb upstream of the closest 5’UTR. Sequences were scanned for motifs using the universalmotif [39] package as described before [40]. In addition to the list of curated motifs [40] all plant transcription factors were obtained from JASPAR [41] and the dicot TATA-box core motif was obtained from PlantProm [42]. Graphs were made with the R packages ggplot2, gggenes, ggbeeswarm, ggpubr and ggtext [43–47] and figure parts were combined with the cowplot package [48]. To estimate statistical significance a two-sided student’s t-test was performed using the package rstatix [49]. In addition to the packages mentioned above the following packages were used: here, R.utils, data.table, readxl, openxlsx, progress, bibtex and several packages within the tidyverse [50–56].

## Results

### Genome search

To find potential novel enhancer sequences, we queried the genomes of different crop and model plant species for the presence of known enhancer sequences (Figure 1, Table 1).

**Figure 1:**
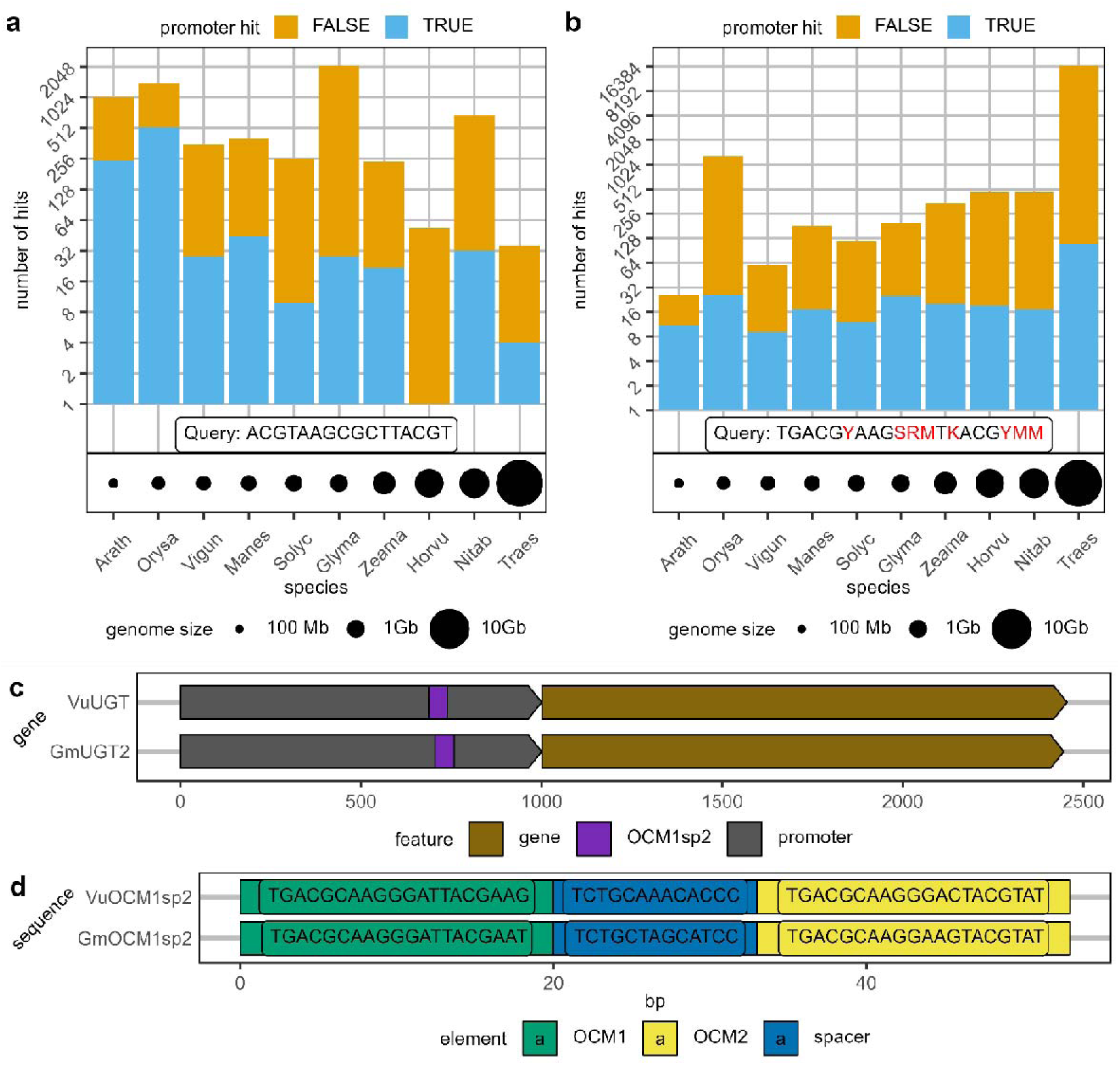
Genome search results. Different plant genomes were queried for the presence of small enhancer-like sequences using BLAST (a) and Biostrings (b), with the 16 bp ocs sequence and the 20 bp ocs consensus sequence respectively. Bars show the number of hits on a logarithmic scale and are colored depending on whether a hit was found in a promoter of a gene (TRUE: blue) or not (FALSE: orange). Query sequence is shown in a box below the graph with degenerate codes colored red. Circles below bar graph represent genome size. c) Schematic gene overview of two orthologous UDP-glucosyl transferase genes from soybean (GmUGT2) and cowpea (VuUGT), with a region that has two OCMs separated by a spacer (OCM1sp2) in their promoter. Promoter comprises the first 1000 bp upstream of the first codon and gene comprises region from first to last codon. d) Schematic overview and nucleotide sequence of OCM1sp2 regions in the respective genes shown in c. Mb: Megabase, Gb: Gigabase, bp: base pair

**Table 1:**
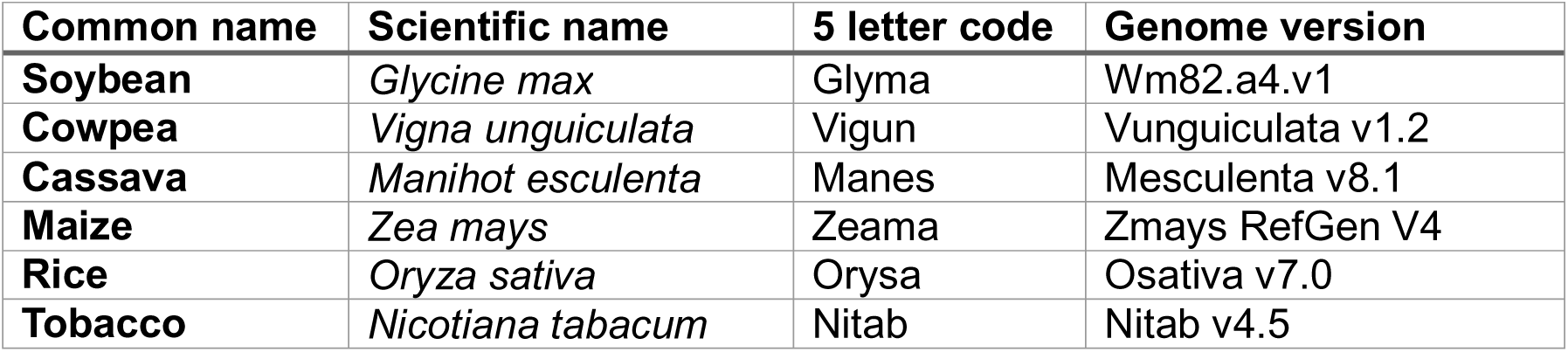
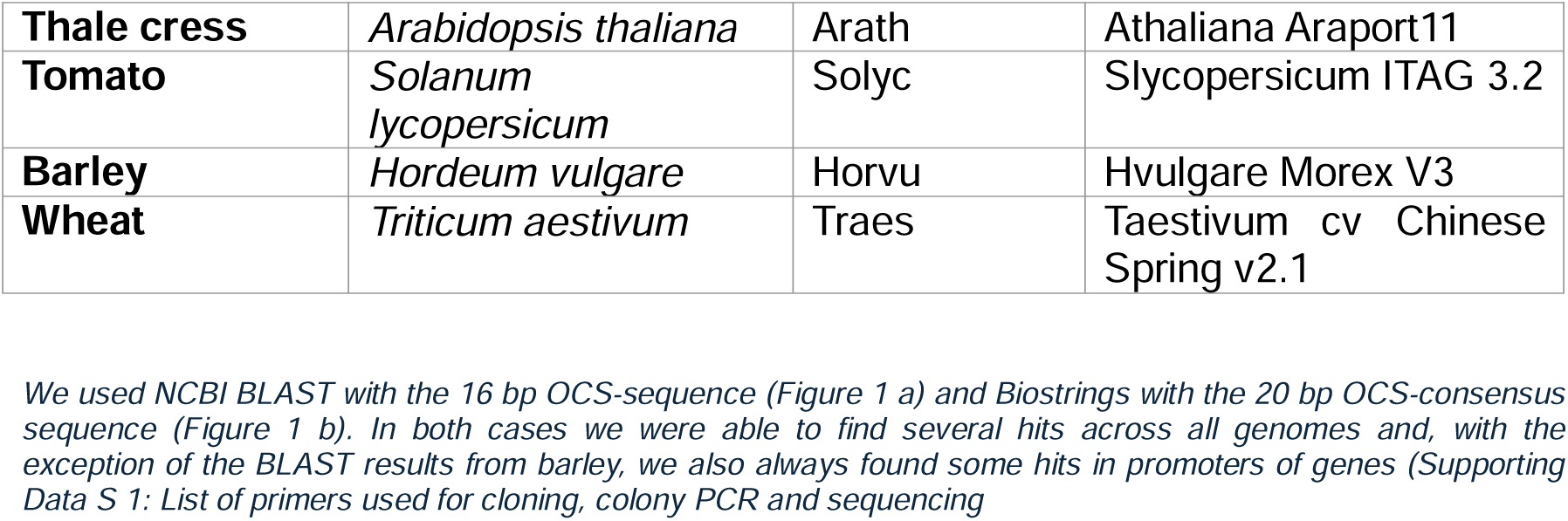
Plant species and respective genomes used for genome query.

Supporting Data S 2, Supporting Data S 3). Perhaps surprisingly, we found a negative trend of a decreasing number of hits with increasing genome size (Figure 1 a). Using Biostrings and the 20 bp consensus sequence we found the opposite trend of an increasing number of hits with an increase in genome size (Figure 1 b), which appears more plausible, especially considering that some query nucleotides are degenerate bases. In a simulated genome, which contains concatenated randomly sampled nucleotides, the number of hits was comparable to genomes of similar size (Supporting Figure S 1). Based on the Biostrings approach we found 25 and 9 hits in the promoters of soybean and cowpea, respectively (Supporting Data S 3). We are referring to these hits as OCS-consensus match (OCM). When taking a closer look at these hits we found the genes GmUGT2 (Glyma.11G000500)[57] and VuUGT (Vigun02g204200) in each of the genomes, which each have two OCMs separated by a small spacer (OCM1sp2)(Figure 1 c + d, Supporting Data S 3). For our downstream validation we derived our first putative enhancer constructs from this motif.

### Fluorescence analysis

As target promoters for the validation experiments, we used the promoters of SBPase in soybean (GmSBP1/Glyma.11G226900 + GmSBP2/Glyma.18G030400) and cowpea (VuSBP/Vigun01g224100). Using a previously published approach [40], we scanned the respective promoters for the presence of known transcription factor binding sites to find a site where an insertion would cause the least disruption (Supporting Data S 4 - Supporting Data S 6). For the first set of enhancers we selected the second OCM in GmUGT2 and VuUGT (OCM2), because this motif has both a “TGACG” motif as well as an “ACGT” motif, which had both previously been hypothesized to be relevant for enhancing gene expression [10, 58, 59]. To achieve a similar length as the published plant enhancer which is a trimer (3xPE; 3×12 bp = 36bp) [7], we designed the OCM2 motifs as dimers (2xOCM2; 2×20 bp = 40 bp). (Figure 2 a). After cloning wild type promoters and constructs where these motifs replaced part of the wild type promoters into our reporter vector, we used the resulting constructs for our transient YFP/RFP-assay and measured relative fluorescence intensity (Figure 2 b, see Supporting Data S 7 for a list of all used enhancer motifs). We found that the 3xPE constructs, induced relative fluorescence intensity significantly. In GmSBP1 the induction was ∼3.31-fold in comparison to the wild type value, in GmSBP2 the induction was ∼1.99-fold and in VuSBP the induction was ∼2.03-fold (Supporting Data S 8). The 2xOCM2 constructs however showed wild type-like fluorescence levels in GmSBP1 and GmSBP2 (Figure 2 b). In VuSBP the 2xOCM2 construct shows a moderate but significant induction, which likely can be attributed to the creation of a TATA box at the end of the insertion. In fact, when testing another construct where we moved the whole motif 1 base pair downstream to disrupt the TATA motif, we did not see a significant increase (Supporting Figure S 2).

**Figure 2:**
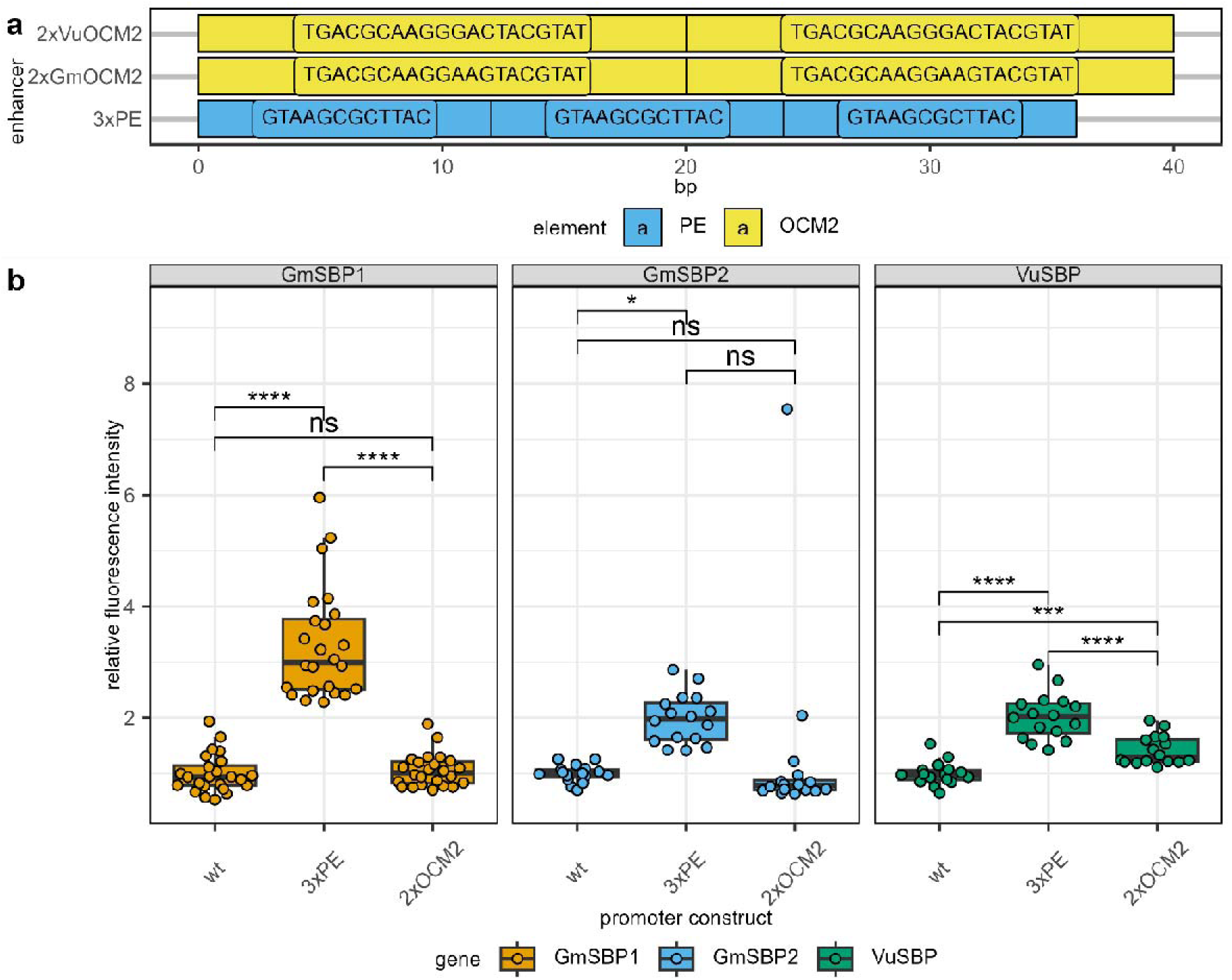
Enhancer sequences and fluorescence values of first set of constructs. a) Schematic overview and sequence of tested enhancers. b) Relative fluorescence intensity of leaf discs punched out of leaf regions, infiltrated with A. tumefaciens carrying constructs with the respective enhancers. Separate plot panels show results from different promoters. Data shown as box- and pointplot with individual points representing values of biological replicates. Boxplot uses box from lower quartile to upper quartile and median value as middle line. Regions outside the box with values deviating up to 1.5 the interquartile range from the quartiles are shown by vertical lines. Values from 2-3 separate experiments were pooled together. Statistical significance was estimated with a two-sided student’s t-test. Brackets indicate compared group and level of significance (*:p< 0.05; **: p<0.01, ***: p<0.001; *: p<0.0001; ns: not significant). N = 16 -24; bp: base pair

For our next set of promoters we tested whether it would need a trimer of the OCM2 to elicit an induction of gene expression and if the OCM1sp2 region as present in the promoters of GmUGT2 and VuUGT would have an induction effect (Figure 3a). Again, we tested the resulting constructs in our transient fluorescence assay (Figure 3 b). We found that the OCM1sp2 construct significantly increased relative fluorescence intensity in all promoters. The induction was in average ∼3.57-fold in GmSBP1, ∼3.03-fold in GmSBP2 and ∼2.15-fold in VuSBP (Supporting Data S 8). For the 3xOCM2 construct we could only see a minor increase in fluorescence intensity in VuSBP, which is likely due to the creation of another TATA-motif.

**Figure 3:**
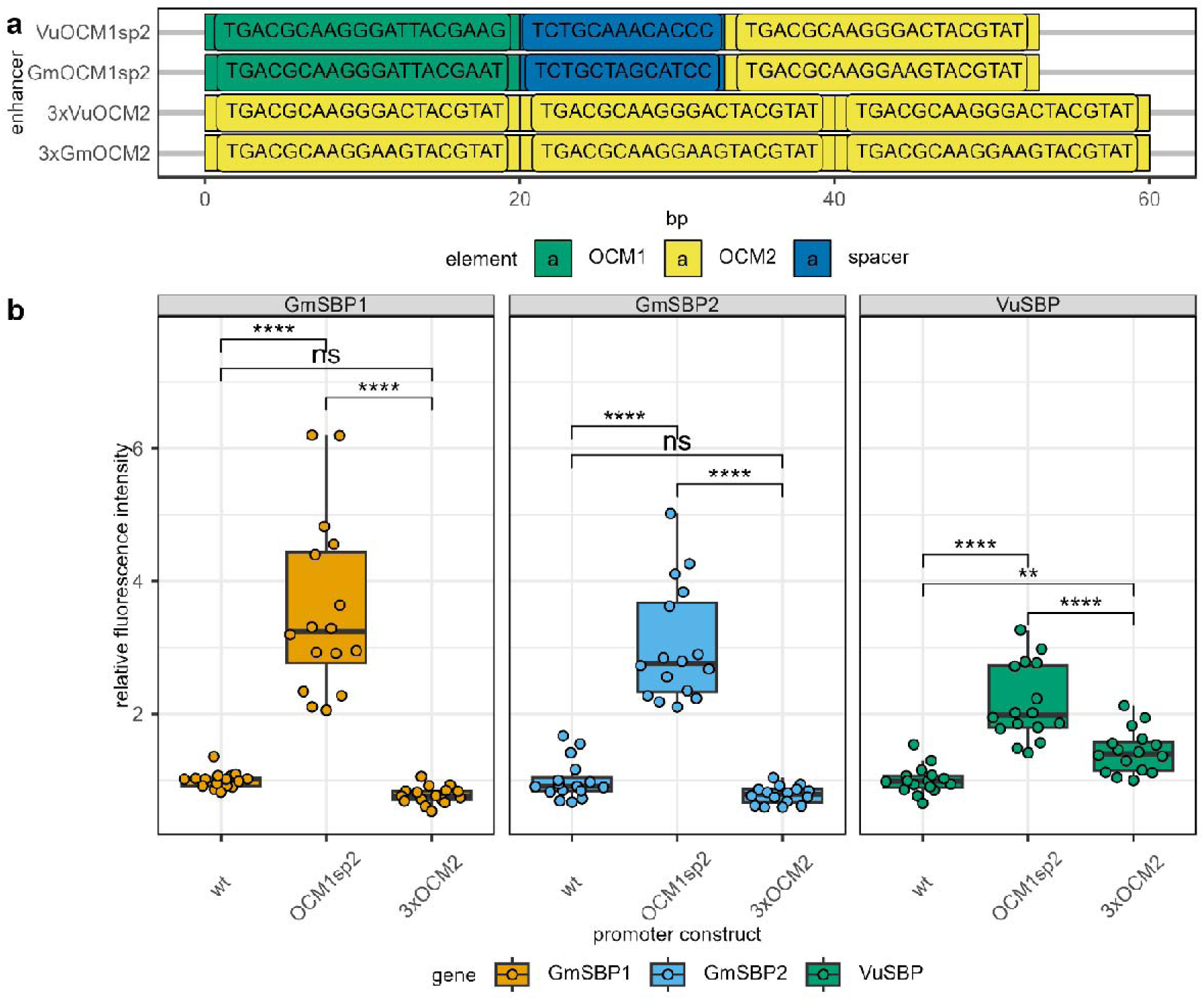
Enhancer sequences and fluorescence values of second set of constructs. a) Schematic overview and sequence of tested enhancers. b) Relative fluorescence intensity of leaf discs punched out of leaf regions, infiltrated with A. tumefaciens carrying constructs with the respective enhancers. Separate plot panels show results from different promoters. Data shown as box- and pointplot with individual points representing values of biological replicates. Boxplot uses box from lower quartile to upper quartile and median value as middle line. Regions outside the box with values deviating up to 1.5 the interquartile range from the quartiles are shown by vertical lines. Values from 2 separate experiments were pooled together. Statistical significance was estimated with a two-sided student’s t-test. Brackets indicate compared group and level of significance (*:p< 0.05; **: p<0.01, ***: p<0.001; *: p<0.0001; ns: not significant). N = 16; bp: base pair

We hypothesized that the inductive activity on the enhancer stemmed from the first OCM, based on a previous observation that a “TGACG” pentamer followed by 7 random bases and another “TGACG” pentamer may be important for the enhancement [10, 58]. Since OCM1 and 2 are both followed by a “C” in soybean and cowpea and we wanted to test whether a “T” at this position is crucial or beneficial, we went back to the genome search to look for such a motif. We found such a motif (OCM3) in the promoter of the genes GmNAC039 (Glyma.06G157400) and VuNAC (Vigun09g119900) in the soybean and cowpea genome (Figure 4 a, Supporting Data S 3). To create the 5 + 7 bp motifs we used both OCM2 and OCM3 and trimmed them to 12 bp from the 3’ end (OCM2t3 + OCM3t3) and then triplicated the 12 bp (Figure 4 a). We used the resulting constructs again for our transient assay and measured relative fluorescence intensity (Figure 4 c). We could see that all enhancer constructs significantly increased relative fluorescence intensity in all promoters in comparison to the respective wild type promoter. For the 3xOCM2t3 constructs, the induction was ∼3.63-fold for GmSBP1, ∼2.05-fold for GmSBP2 and ∼3.08-fold for VuSBP (Supporting Data S 8). For the 3xOCM3t3 constructs the induction was ∼4.28-fold for GmSBP1, ∼2.53-fold for GmSBP2 and ∼3.43-fold for VuSBP (Supporting Data S 8). In comparison between the two constructs the OCM3t3 showed a small but significant higher induction than OCM2t3 in GmSBP1 and GmSBP2.

**Figure 4:**
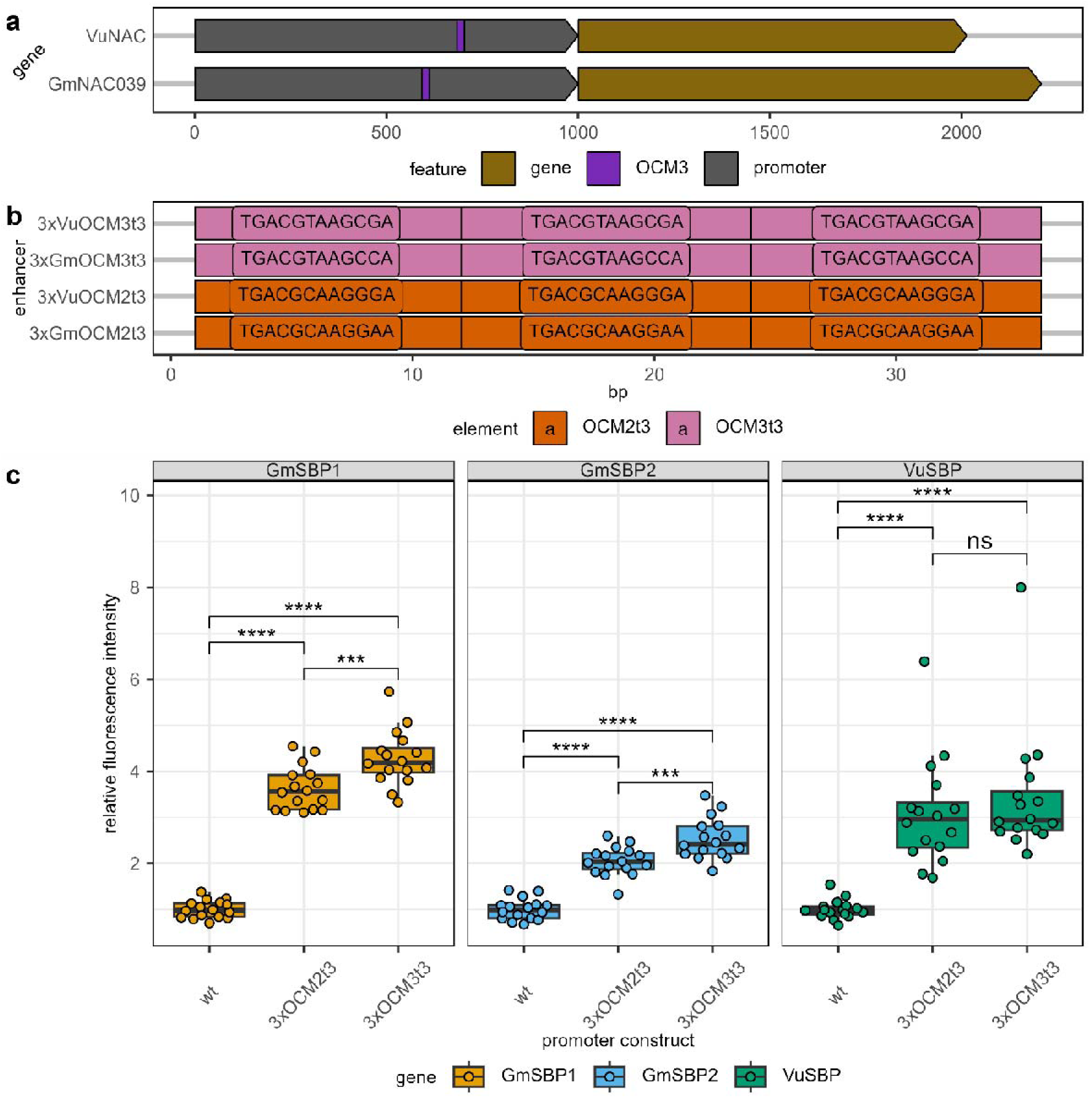
Enhancer sequences and fluorescence values of third set of constructs. a) Schematic gene overview of two orthologous NAC genes from soybean (GmNAC039) and cowpea (VuNAC), with a region that has an OCM (OCM3) in their promoter. Promoter comprises the first 1000 bp upstream of the first codon and gene comprises region from first to last codon. b) Schematic overview and sequence of tested enhancers. c) Relative fluorescence intensity of leaf discs punched out of leaf regions, infiltrated with A. tumefaciens carrying constructs with the respective enhancers. Separate plot panels show results from different promoters. Data shown as box-and pointplot with individual points representing values of biological replicates. Boxplot uses box from lower quartile to upper quartile and median value as middle line. Regions outside the box with values deviating up to 1.5 the interquartile range from the quartiles are shown by vertical lines. Values from 2 separate experiments were pooled together. Statistical significance was estimated with a two-sided student’s t-test. Brackets indicate compared group and level of significance (*:p< 0.05; **: p<0.01, ***: p<0.001; *: p<0.0001; ns: not significant). N = 16; bp: base pair

Because we saw differences in the fold-induction of the different constructs, in different promoters, we looked at the unnormalized base-level relative fluorescence intensity of the wild type promoters (Figure 5). The mean values of relative fluorescence intensity were ∼2.89 for GmSBP1, ∼5.63 for GmSBP2 and ∼3.08 for VuSBP (Supporting Data S 9). The expression level in GmSBP2 was thus significantly higher than the one of GmSBP1 and VuSBP (Figure 5).

**Figure 5:**
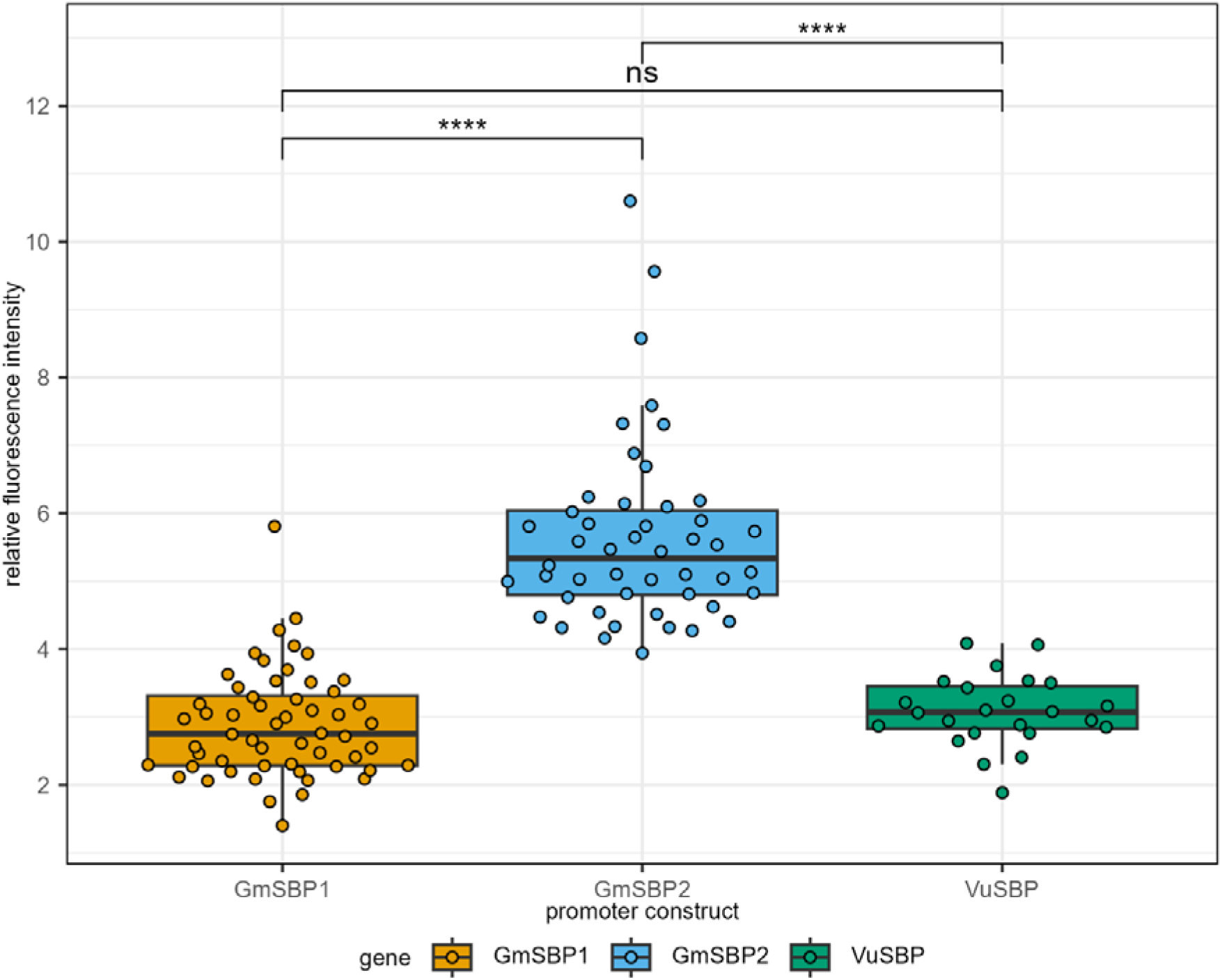
Fluorescence values of wild type promoters. Relative fluorescence intensity of leaf discs punched out of leaf regions, infiltrated with A. tumefaciens carrying constructs with the respective construct. Data shown as box-and pointplot with individual points representing values of biological replicates. Boxplot uses box from lower quartile to upper quartile and median value as middle line. Regions outside the box with values deviating up to 1.5 the interquartile range from the quartiles are shown by vertical lines. Values from all experiments were pooled together. Statistical significance was estimated with a two-sided student’s t-test. Brackets indicate compared group and level of significance (*:p< 0.05; **: p<0.01, ***: p<0.001; *: p<0.0001; ns: not significant). N = 16 -58

Finally, we were interested to see if any known transcription factors might be able to bind to the enhancer sequences we tested. To this end we performed another motif scan on the final promoter construct sequences, where we scanned the 1000 bp upstream of the first YFP codon for the presence of any motifs (Figure 6, Supporting Data S 10). In constructs containing the 3xPE enhancer we found two sites, with partially overlapping binding sites for TGA and NAC transcription factors. In both GmSBP2 and VuSBP we found another site for NAC transcription factors at the start of the enhancer. In the 2xOCM2 constructs we found two overlapping sites for SPL4 and Zm00001d018571 in GmSBP1 and GmSBP2 and two sites for TCP5 in VuSBP (Figure 6, Supporting Data S 10). Similarly, we found one such site at the end of OCM1sp2 constructs and 3 such sites in 3xOCM2 constructs. In GmSBP1 we additionally found some sites for NAC transcription factors, as well as AT1G19040 at the start of the enhancer (Figure 6, Supporting Data S 10). Surprisingly, we found almost no motifs in the constructs with 3xOCM2t3, that are not also in the wild type promoter (Figure 6, Supporting Data S 10). The only exception is a site for a myb-related transcription factor cluster at the end of the enhancer of VuSBP. Contrary, in the constructs with the 3xOCM3t3 we found three sites in all promoters where different TGA and bZIP transcription factors could bind (Figure 6, Supporting Data S 10).

**Figure 6:**
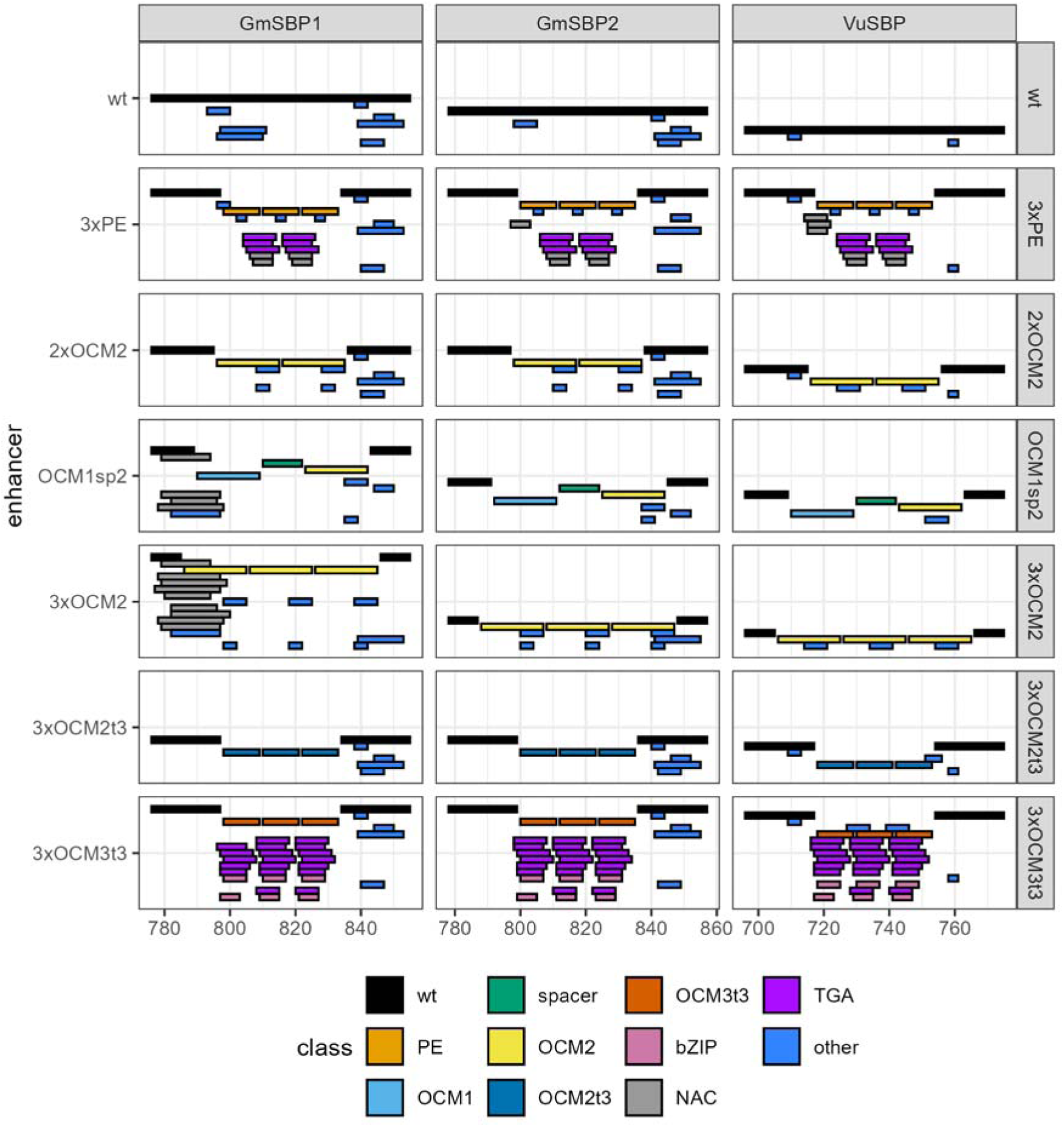
Putative motifs in wild type promoters and promoters with insertion of enhancer motif. Each panel shows an individual promoter construct. Plots are facetted into columns by promoter and rows by enhancer type. X-axis shows the position within the promoter i.e. the 1000 bp before the first codon of EYFP. For better visibility only the regions around the enhancer insertion are shown and only motifs are displayed that do not occur in all constructs of the respective promoter. Motifs and enhancer elements are shown as color-filled rectangles and spread across the Y-axis to allow distinguishing overlapping motifs.

## Discussion

In this study we found hits for sequences resembling the OCS sequence in the genomes of different plant species both based on the original OCS sequence as well as the OCS consensus sequence [8, 10]. Using two separate approaches we found different numbers of hits, which is likely to be explained by the difference in the search algorithm and length and identity of the query sequence. Several of these hits resided in promoter sequences of genes. We selected a few of these and tested derivatives of these sequences in our transient assay. We could find enhancing effects for the 3xPE, OCM1sp2, 3xOCM2t3 and 3xOCM3t3 constructs. For the 3xPE and the 3xOCM3t3 constructs we could further find that the insertion of these motifs created binding sites for TGA-transcription factors and other factors. It therefore seems likely that enhancement of gene expression is achieved by transcription factor binding. Even though we could not find these TGA-TF binding sites in OCM1sp2 and 3xOCM2t3, with the used motif scan, it seems plausible, that similar factors may bind to these motifs. As already shown by early experiments, binding of two units is likely necessary for transcriptional enhancement [9]. This is also in agreement with findings documented in the patent (US11198885B1) related to the 3xPE [7]. There they found that a 2xPE is not sufficient to induce gene expression enhancement, which based on our results of the promoter scan should only create one binding site for TGA-like factors. In the case of the full OCM2 constructs (2xOCM2 + 3xOCM2) it remains unclear, why these did not lead to a gene expression enhancement but based on previous results we can have two hypotheses. The first explanation would be that the OCM2 element itself is not a functional enhancer, in which case the enhancement in the OCM1sp2 construct is exclusively mediated by OCM1. Focusing on TGA-TF-like binding sites in these motifs, both of these motifs start with a “TGACG” motif, followed by 7 bases. In the case of OCM1 this is followed by “TTACG” and in the case of OCM2 this is followed by “CTACG” and it could be possible that the latter does not have a high enough affinity for TGA-like factors. It could previously be shown that individual mutations at specific sites of the 16 bp OCS element can have strong effects on the enhancement capability of the motif [9]. The second explanation could be that the stacking of the 20 bp motifs in the full OCM2 constructs did not create the appropriate spacing of potential binding sites. Binding of TGA-factors to the first TGA-motif of a OCM2 motif could prevent binding of the second TGA-motif in the previous OCM2. Correct spacing of motifs has been suggested to be crucial for transcriptional enhancement as both insertions and deletions can remove the enhancing effect of OCS-elements [9, 59]. In this case it could be possible that both OCMs in OCM1sp2 constructs are able to bind TGA-like factors.

Several-follow up experiments could be performed to further characterize the enhancer motifs. The number of multimers of the motifs could be both increased and decreased. By adding more motifs, it may be possible to further increase the enhancing effect of the sequences and it could thus be tested how many motifs can be stacked before gene expression does not further increase. As shown in the patent (US11198885B1, Figure 2) related to the 3xPE [7] adding a tetramer (4x) even lead to a decrease again in comparison to the trimer (3x) in the promoter of ZmGln1-3. It could be possible that the enhancing effect of the sequences also depends on the specific promoter. In our experiments we did find differences in the induction capability of enhancers in the context of different promoters, which may in turn be related to their baseline expression levels. This is consistent with results from the 3xPE, where gene expression could be enhanced in almost all promoters through the insertion of the plant enhancer, but fold-induction diminished towards a higher baseline expression [7] (see Figure 1 e in [7]). Reducing the amount of motifs would also be interesting, in order to find out what the minimal sequence is that is required for enhancement of gene expression. As mentioned before, based on previous findings we can expect two binding sites to be required for the enhancement effect [9]. Based on our results and predictions, that means a minimal sequence could consist of two “TGACG” motifs with 7 base pairs in between, which would add up to a total of only 17 base pairs. This may be helpful as shorter sequences may be easier to insert into plant promoters by means of gene-editing.

Even though further investigation is required for a mechanistic understanding, our results already show several motifs that can induce target gene expression, which could be used in a biotechnological application. We show that motifs related to known enhancer sequences like the OCS-element can be found in many different plant genomes. As such our approach to find endogenous enhancer sequences and use them to increase the expression of target genes, may be applicable to a large number of target genes and species. As techniques like prime-editing are becoming more feasible in dicot plants [60], our approach may open the door for precise gene-editing in the form of direct insertion of enhancers into promoter sequences. Importantly, as these sequences would be derived from the genome of the respective target plant, such edits may be considered cisgenic.

## Data Availability Statement

All data and workflows related to the findings presented here are managed in an Annotated Research Context in the DataHUB of DataPLANT and will be released upon acceptance of the manuscript for publication.

## Author Contributions

CAR acquired funding for this work. CAR and VM conceptualized the work. MWA performed investigation, formal analysis, visualization and wrote the original manuscript draft. All authors were involved in reviewing and editing the manuscript.

## Acknowledgments

This work was supported by the project Realizing Increased Photosynthetic Efficiency (RIPE), that is funded by Gates Agricultural Innovations grant investment 57248, awarded to the University of Essex, UK by the University of Illinois, USA. The authors would like to acknowledge the use of the High Performance Computing Facility (Ceres) for this work and thank Stuart Newman for support in using the facility at the University of Essex. We would also like to thank Professor Phillip Mullineaux for support with the fluorescence assays on the plate reader and for supplying an RFP vector sequence.

## Supporting information

*Supporting Data S 1: List of primers used for cloning, colony PCR and sequencing*

*Supporting Data S 2: Genome hits as found by NCBI BLAST*

*Supporting Data S 3: Genome hits as found by Biostrings*

*Supporting Data S 4: Potential transcription factor binding sites in GmSBP1 found by promoter motif scan*

*Supporting Data S 5: Potential transcription factor binding sites in GmSBP2 found by promoter motif scan*

*Supporting Data S 6: Potential transcription factor binding sites in VuSBP found by promoter motif scan*

*Supporting Data S 7: Sequence list of used enhancer motifs*

*Supporting Data S 8: Average normalized fluorescence values of all constructs and assays*

*Supporting Data S 9: Average relative fluorescence intensity of wild type promoter constructs across all assays*

*Supporting Data S 10: Potential transcription factor binding sites in used promoter constructs*

*Supporting Figure S 1: Genome search results using Biostrings and the 20 bp ocs consensus sequence*.

*Supporting Figure S 2: Fluorescence values from VuSBP promoter constructs with 2xOCM2 enhancer moved 1 base pair downstream*.

