## Supporting Figure for "Endogenous short enhancer sequences increase expression of soybean and cowpea RUBP regeneration genes"

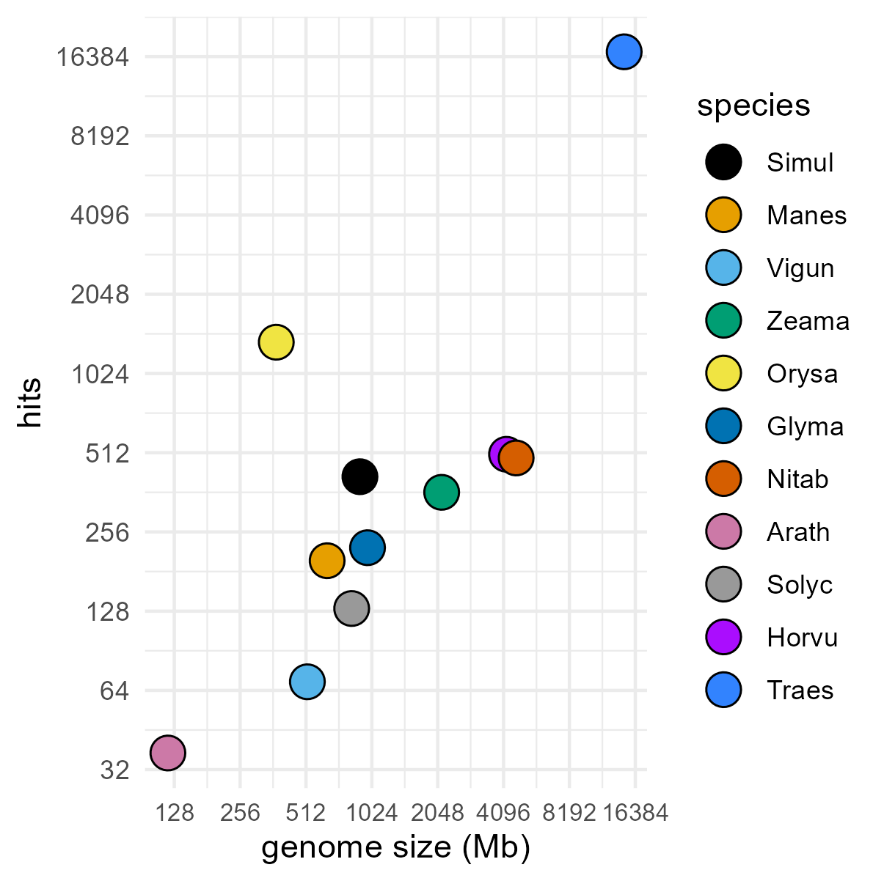

Supporting Figure S 1: Genome search results using Biostrings and the 20 bp ocs consensus sequence. Different plant genomes were queried for the presence of small enhancer-like sequences using Biostrings with the 20 bp ocs consensus sequence. X-axis shows genome size of respective plant genome and Y-axis shows the number of hits. Both axes use logarithmic scale. Each color-filled dot represents a different genome. Mb: Megabase

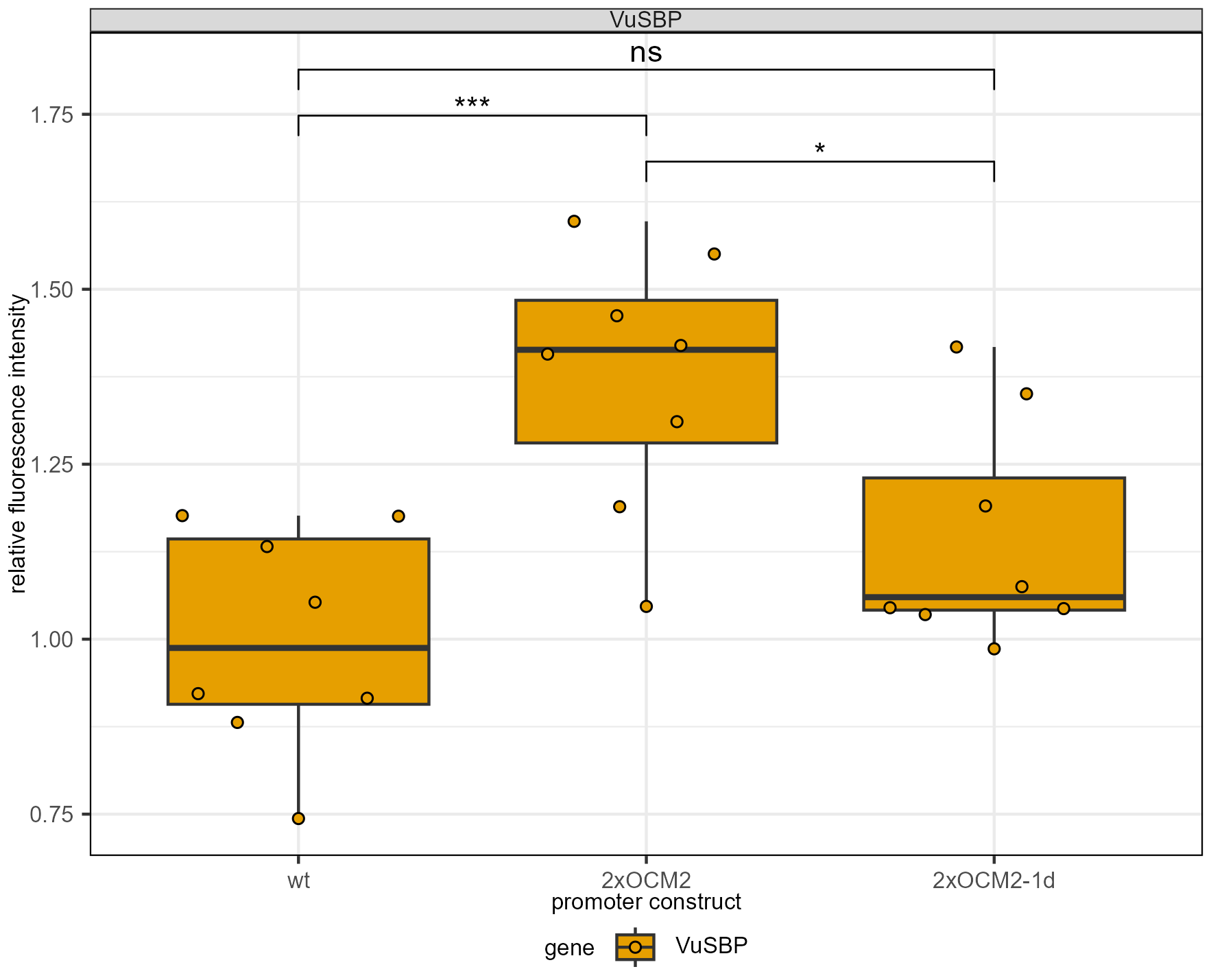

Supporting Figure S 2: Fluorescence values from VuSBP promoter constructs with 2xOCM2 enhancer moved 1 base pair downstream. Relative fluorescence intensity of leaf discs punched out of leaf regions, infiltrated with A. tumefaciens carrying constructs with the respective enhancers. Data shown as boxplot overlaid with individual points representing values of biological replicates. Boxplot uses box from lower quartile to upper quartile and median value as middle line. Regions outside the box with values deviating up to 1.5 the interquartile range from the quartiles are shown by vertical lines. Values from 2-3 separate experiments were pooled together. Statistical significance was estimated with a two-sided student’s t-test. Brackets indicate compared group and level of significance (*:p< 0.05; **: p<0.01, ***: p<0.001; *: p<0.0001; ns: not significant). N = 8
